# Animal models with group-specific additive genetic variances: extending genetic group models

**DOI:** 10.1101/331157

**Authors:** Stefanie Muff, Alina K. Niskanen, Dilan Saatoglu, Lukas F. Keller, Henrik Jensen

**Affiliations:** Institute of Evolutionary Biology and Environmental Studies, University of Zurich, Winterthurerstrasse 190, Zurich, Switzerland; Epidemiology, Biostatistics and Prevention Institute, Department of Biostatistics, University of Zurich, Hirschengraben 84, Zurich, Switzerland; Centre for Biodiversity Dynamics, Department of Biology, Norwegian University of Science and Technology, Høgskoleringen 5, Trondheim, Norway; Department of Ecology and Genetics, University of Oulu, P.O. Box 3000, Oulu, Finland; Zoological Museum, University of Zurich, Karl-Schmid-Strasse 4, Zurich, Switzerland

**Keywords:** Relatedness matrix, immigration, Cholesky decomposition, partial inbreeding coefficient, house sparrows

## Abstract

**1.** The *animal model* is a key tool in quantitative genetics and has been used extensively to estimate fundamental parameters, such as additive genetic variance, heritability, or inbreeding effects. An implicit assumption of animal models is that all founder individuals derive from a single population. This assumption is commonly violated, for instance in cross-bred livestock breeds, when an observed population receive immigrants, or when a meta-population is split into genetically differentiated subpopulations. Ignoring genetic differences among different source populations of founders may lead to biased parameter estimates, in particular for the additive genetic variance.

**2.** To avoid such biases, genetic group models, extensions to the animal model that account for the presence of more than one genetic group, have been proposed. As a key limitation, the method to date only allows that the breeding values differ in their means, but not in their variances among the groups. Methodology previously proposed to account for group-specific variances included terms for segregation variance, which rendered the models infeasibly complex for application to most real study systems.

**3.** Here we explain why segregation variances are often negligible when analyzing the complex polygenic traits that are frequently the focus of evolutionary ecologists and animal breeders. Based on this we suggest an extension of the animal model that permits estimation of group-specific additive genetic variances. This is achieved by employing group-specific relatedness matrices for the breeding value components attributable to different genetic groups. We derive these matrices by decomposing the full relatedness matrix via the generalized Cholesky decomposition, and by scaling the respective matrix components for each group. To this end, we propose a computationally convenient approximation for the matrix component that encodes for the Mendelian sampling variance. Although convenient, this approximation is not critical.

**4.** Simulations and an example from an insular meta-population of house sparrows in Norway with three genetic groups illustrate that the method is successful in estimating group-specific additive genetic variances and that segregation variances are indeed negligible in the empirical example.

**5.** Quantifying differences in additive genetic variance within and among populations is of major biological interest in ecology, evolution, and animal and plant breeding. The proposed method allows to estimate such differences for subpopulations that form a connected meta-population, which may also be useful to study temporal or spatial variation of additive genetic variance.

## 1. Introduction

Quantifying the (causal) relationships between genes and observed phenotypic traits is a central task of empirical studies of adaptive evolution (Lynch & Walsh, 1998; Charmantier *et al.*, 2014) and of plant and animal breeding (Falconer & Mackay, 1996). The *animal model* (Henderson, 1984; Kruuk, 2004; Wilson *et al.*, 2010) has become a popular statistical approach to disentangle genetic effects on a phenotype from other factors that may induce phenotypic similarities among relatives, such as shared environmental effects (Stopher *et al.*, 2012), inbreeding (Reid & Keller, 2010), or individual traits such as age or sex (Wilson, 2008; de Villemereuil *et al.*, 2017). Fundamental to the animal model is information on how animals are related to each other, information typically obtained from pedigree data (Wright, 1922; Henderson, 1976) or from single-nucleotide polymorphism (SNP) data (*e. g*. Speed & Balding, 2015). Pedigrees are still the most commonly used source of relatedness information in animal models (*e. g*. Bonnet *et al.*, 2017; Wolak & Reid, 2017), in part because this leads to models that are computationally efficient.

All pedigrees start with a founder generation with unknown parents (Wolak & Reid, 2017, Appendix S1). The animal model assumes that all founder individuals stem from a single, genetically homogeneous baseline population. When this assumption is violated, for example in populations that receive immigrants or in crossbred livestock breeds, estimates of additive genetic variance are biased (Dong *et al.*, 1988; Cantet *et al.*, 2000). To address this problem, animal breeders developed *genetic group models* (*e. g*. Quaas, 1988), models that are now also receiving attention in evolutionary ecology (Wolak & Reid, 2017). The main idea behind genetic group models is that estimates of additive genetic variance may be biased if animals of distinct genetic origin differ in their mean breeding values, and that accounting for such differences in breeding values may reduce or eliminate the bias (Wolak & Reid, 2017). However, current genetic group models have an important key limitation: genetic groups are allowed to differ in mean breeding value but are assumed to have the same additive genetic variance. This homogeneity assumption is violated in some animal breeding applications (*e. g*. Elzo, 1990; Lo *et al.*, 1993; Alfonso & Estany, 1999), and is likely also violated in many natural populations, where source populations of immigrants may differ in additive genetic variance due to differences in effective population size, selection regimes, etc. (*e. g*. Mackay, 1981; Lande, 1988; Hoffmann *et al.*, 2017).

Aiming for better predictions of breeding values in crossbred populations, animal breeders have suggested approaches that account for heterogeneous additive genetic variances across genetic groups (Lo *et al.*, 1993; Cantet & Fernando, 1995; García-Cortés & Toro, 2006). One drawback of these models is that they rapidly become infeasibly complex, mainly due to additional terms that are needed to account for segregation variance when breeds are mixed. Segregation variance refers to differences in variance when the average proportion of genetic origin from the ancestral genetic groups is the same, as in *F*_1_ and *F*_2_ generations of line crosses (Wright, 1968; Lande, 1981). Segregation variances can be large in crossbreeding applications (Lynch & Walsh, 1998, p. 11). However, as we detail below, we expect the segregation variance from crossing different genetic groups in wild study populations to be small for many traits of interest, so that omitting it from the animal model does not lead to significant bias.

Here we use this simplification to derive genetic group models that allow for group-specific additive genetic variances. Similar to García-Cortés & Toro (2006), we additively split the total breeding value of each individual into group-specific components, which covary according to a group-specific relatedness matrix. The main challenge is to find these matrices, and we derive them here by decomposing the full relatedness matrix via a generalized Cholesky decomposition (as described by Henderson, 1976), and by appropriately scaling the respective matrix components for each group. In the following, we first summarize the current state of the homogeneous genetic group models and then give a detailed description of our extension to heterogeneous group-specific additive genetic variances. We illustrate the performance of our method with a simulation study and an application to a meta-population of house sparrows (*Passer domesticus*) in Norway, where genetic groups are determined by geographical properties of the bird’s natal island population. By also fitting a model that includes the segregation variance, we use this empirical study system to illustrate that omitting segregation variances is unproblematic in such applications. Finally, we discuss opportunities and limitations of our extended genetic group model.

## 2 The animal model for genetic groups with homogeneous variances

The basic animal model for a phenotypic measurement *y*_*i*_ of an individual *i* (1 ≤ *i* ≤ *n*) is

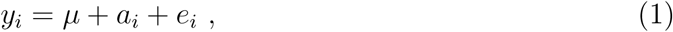

with population mean *µ*, environmental component 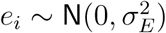 with environmental variance 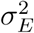, and breeding values distributed as 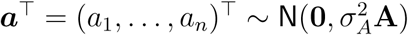 with additive genetic variance 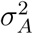 and additive genetic relatedness matrix **A** that represents the relatedness among individuals (Henderson, 1976; Kruuk, 2004). In diploid noninbred organisms, each element in **A** is equal to two times the coefficient of coancestry between a pair of individuals (Lynch and Walsh 1998). Model (1) is often extended by fixed effects (such as sex or age) and by additional random effects that account for permanent environmental conditions (see *e. g*. Wilson *et al.*, 2010). In all cases, the underlying assumption is that all animals in the analysis derive from the same genetic population, and that the breeding values (*a*_*i*_) encode for the deviation from the mean of this population and thus have a mean of zero.

As noted by animal breeders a long time ago, this assumption is frequently violated, for example in crossbred populations from genetically differentiated breeds. In such cases it is necessary to allow for differences in mean breeding values among animals with different genetic origin (Dong *et al.*, 1988; Quaas, 1988). Let us denote by a founder population a set of animals with unknown (*i. e*. missing) parents, and assume that *r* founder populations exist, where each of them corresponds to a different genetic group. When animals from different genetic groups mate, their genetic contributions are propagated through the pedigree following the Mendelian rules of inheritance. Offspring in later generations may thus inherit different proportions of the genome from the genetic groups. Denote by *q*_*ij*_ the expected proportional contribution of founder population *j* to the genome of individual *i*. The respective values are typically written into a matrix **Q** with *n* rows (*n*=number of animals) and *r* columns, such that *q*_*ij*_ is the value in the *i*^*th*^ row and *j*^*th*^ column (Wolak & Reid, 2017, Fig. 3). Following the notation of Wolak & Reid (2017), and denoting by *g*_*j*_ the expected average genetic effect in group *j*, the basic animal model (1) is then extended to

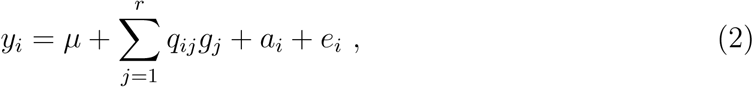

with total additive genetic effect of individual *i* given as 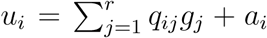, that is, the weighted sum of genetic group-mean effects, plus the breeding value *a*_*i*_ of the individual that accounts for deviations from the weighted group mean. This model allows for different mean, but it is actually overparameterized because for each *i* the contributions from the *r* groups sum up to 1, that is, 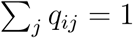. Similar to ANOVA models or when categorical variables are included in regression models, the parameters become identifiable when one group is set as reference group (*e. g*. assuming *g*_1_ = 0), or when additional constraints are added, such as 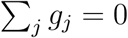.

Let us illustrate the idea for two genetic groups. When using the convenient constraint *g*_1_ = 0, phenotypes of animals that have ancestors either only from group 1 (*i. e*. *q*_*i*1_ = 1*, q*_*i*2_ = 0) or only from group 2 (*i. e*. *q*_*i*1_ = 0*, q*_*i*2_ = 1) can be described by the following models

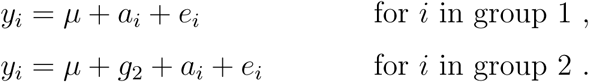

Thus, members of genetic group 1 have a total additive genetic effect of *u*_*i*_ = *a*_*i*_ and members of genetic group 2 have *u*_*i*_ = *g*_2_ + *a*_*i*_, where *g*_2_ estimates the difference between the mean breeding values of the groups. Therefore, the respective values are distributed around an overall mean of 0 and *g*_2_, respectively, but with the same additive genetic variance 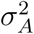 and relatedness matrix **A**. Note that, while the main benefit of including group-specific means is that bias in the estimates of 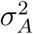 is reduced, the estimated values (*e. g*. of *g*_2_) may sometimes be of interest themselves (see the Conclusions in Wolak & Reid, 2017).

## 3. Genetic group models with heterogeneous additive genetic variances

### 3.1 Segregation variance for polygenic traits

The key limitation of the genetic group model (2) is that all genetic groups are assumed to have the same additive genetic variance 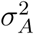, and animal breeders have therefore suggested extensions that allow 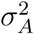 to differ among groups (*e. g*. Lo *et al.*, 1993; Cantet & Fernando, 1995; García-Cortés & Toro, 2006). However, these methods quickly become computationally demanding because the respective models include terms to account for the segregation variance between any two genetic groups, thus *g*(*g* − 1)/2 segregation variances in the presence of *g* genetic groups. The magnitude of these variances may be considerable in artificial breeding scenarios, *e. g*. when crossing genetically differentiated pure-bred lines (see *e. g*. Lande, 1981, Table 3), and if a trait is determined by one or only a few loci. To understand why, let us start by looking at a hypothetical trait that is determined by *m* loci. Following Lynch & Walsh (1998, equation 9.15), the segregation variance between two genetic groups (*e. g*. breeds) can be computed as

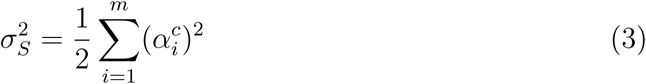

where 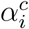 denotes the mean additive genetic difference between the groups due to locus *i*. Let us first look at an extreme example where only one locus determines a trait value, and where two populations or breeds differ in their mean breeding value by *α*^*c*^ = 1 (which is the range of what we will find in the house sparrows example of Section 5). The segregation variance expected in a cross between these breeds is then given as 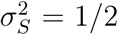 · 1 = 0.5. Genome-wide association studies (GWAS) suggest however that complex (continuous) traits are mostly polygenic, thus the additive genetic component is not determined by a single locus (*e. g*. Goddard & Hayes, 2009; Flint & Mackay, 2009; Hill & Kirpatrick, 2010; Yang *et al.*, 2011b; Robinson *et al.*, 2013; Santure *et al.*, 2015; Silva *et al.*, 2017; Bouwman *et al.*, 2018). In fact, it is a fundamental assumption of quantitative genetics that phenotypic traits are determined by many genes that each contribute a small effect to trait variation, known as the “infinitesimal model” (Fisher, 1918; Bulmer, 1971). If, for example, 100 loci each contribute the same proportion of 1/100 to the overall group difference of 1 in the above example, the segregation variance reduces to

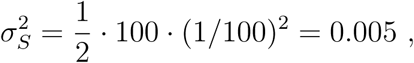

which is exactly 1/100 of the segregational variance for a single-locus trait. Thus for any number of loci *m*, the segregation variance is 1/*m* of what it would be for a single locus, given that each locus contributes the same proportion of the effect. Even when considering that the locus-specific effect sizes are typically heterogeneous, the most influential loci often only explain a small proportion of the phenotypic variance (Yang *et al.*, 2011b; Wood *et al.*, 2014; Husby *et al.*, 2015). Thus, the segregation variance 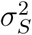 for complex continuous traits is expected to be small compared to the total phenotypic variance in many study systems. In the following extension of the animal model we therefore ignore the segregation variance.

### 3.2 Animal model for heterogeneous additive genetic variances

To allow for heterogeneous additive genetic variances in the animal model, we extend model (2) by splitting the breeding value *a*_*i*_ into group-specific contributions (similar to García-Cortés & Toro, 2006), such that

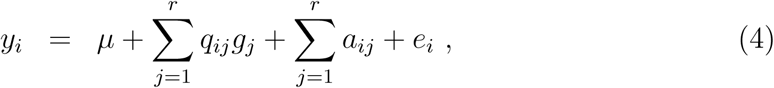

with 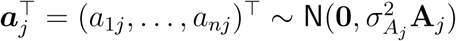 for all groups *j* = 1*, …, r*, where 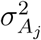 is the additive genetic variance in group *j*, and **A**_*j*_ is a group-specific relatedness matrix. The contribution *a*_*ij*_ to the breeding value of individual *i* can be interpreted as the part that is inherited from group *j*. We assume that the contributions *a*_*ij*_ are independent of each other, because they differ in genetic origin.

We illustrate the idea again for the case of two genetic groups. As above, we constrain the mean breeding value of group 1 (the reference group) to *g*_1_ = 0 for identifiability reasons. In addition, the model must ensure that a breeding value component is zero (*a*_*ij*_ = 0) if the animal’s genome obtains no contribution from the respective group *j* (thus if *q*_*ij*_ = 0), which will be ensured by appropriate choice of the covariance matrix **A**_*j*_ (details will follow in Section 3.3). Animals in genetic group 1 (*i. e*. *q*_*i*1_ = 1) then have a genetic effect *u*_*i*_ = *a*_*i*1_ and animals in genetic group 2 (*i. e*. *q*_*i*2_ = 1) receive *u*_*i*_ = *g*_2_ + *a*_*i*2_, and the breeding values are distributed as

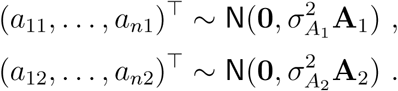

While it is relatively straightforward to formulate such a model, it is less obvious what the group-specific relatedness matrices **A**_*j*_ are. It is, for example, not valid to use **A** for **A**_*j*_, because the within-group relatedness structure is different from the overall relatedness. In addition, the total breeding value is now split into the sum Σ_*j*_ *a*_*ij*_, with *a*_*ij*_ equal to the total breeding value only if an animal has *q*_*ij*_ = 1 for group *j*. The entries of **A**_*j*_ must therefore be scaled accordingly. We now turn to this issue.

### 3.3 Group-specific relatedness matrices

#### 3.3.1 Decomposition of the relatedness matrix

To understand how to specify the group-specific relatedness matrices **A**_*j*_, let us first recall a result by Henderson (1976) for efficient calculation of inverse relatedness matrices. He suggested to decompose **A** by a *generalized Cholesky decomposition* into

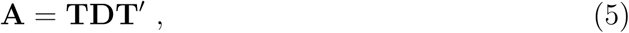

where **T** is lower triangular matrix with transposed **T**′, and **D** = Diag(*d*_11_*, …, d*_*nn*_) is a diagonal matrix with entries *d*_11_*, …, d*_*nn*_. This is equivalent to the Cholesky decomposition **A** = **LL**′ for a lower triangular matrix 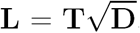. A useful property of the decomposition (5) is that the matrices **T** and **D** have elegant interpretations: **T** traces the flow of alleles from one generation to the other, and the diagonal entries of **D** scale the Mendelian sampling variance (Mrode, 2005, p. 27).

Let us illustrate these properties with an example that we adapted from Mrode (2005, Table 2.1), without genetic groups. The pedigree is given in Table 1, with a corresponding graphical representation of parent-offspring relations (Figure 1a) and matrices **A**, **T** and **D** (Figures 1 b-d), where the generalized Cholesky decomposition to obtain **T** and **D** was calculated with the function `gchol()` that is available in the `bdsmatrix` package (Therneau, 2014) in R (R Core Team, 2017), as pointed out *e. g*. by Gorjanc (2011). In this example, animals 1, 2 and 3 have unknown parents and are denoted as *founders* of the population. Each off-diagonal entry in **T** corresponds to the relatedness coefficient (expected relatedness) of individuals with their direct descendants (*i. e*. children, grandchildren etc.), where columns represent ancestors and rows descendants. For example, individual 1 is the parent of animals 4 and 5, thus the entries (4,1) and (5,1) in the matrix are 0.5. In addition, animal 6 is the offspring of animals 4 and 5, thus the relatedness of 1 with 6 is also 0.5. Finally, the relatedness of 1 and 7 is 0.25. These considerations can be repeated for each column in the matrix, where all diagonal elements are 1 and all elements below the diagonal in the respective column correspond to the *expected* proportion of the genome that is transmitted from the respective ancestor to its direct descendants.

**Table 1:** Pedigree example, adapted from Mrode (2005, Table 2.1).

| ID | Dam | Sire |
| --- | --- | --- |
| 1 ($g_1$) | NA | NA |
| 2 ($g_1$) | NA | NA |
| 3 ($g_2$) | NA | NA |
| 4 | 1 | 2 |
| 5 | 1 | 3 |
| 6 | 5 | 4 |
| 7 | 6 | 3 |

**Figure 1:**
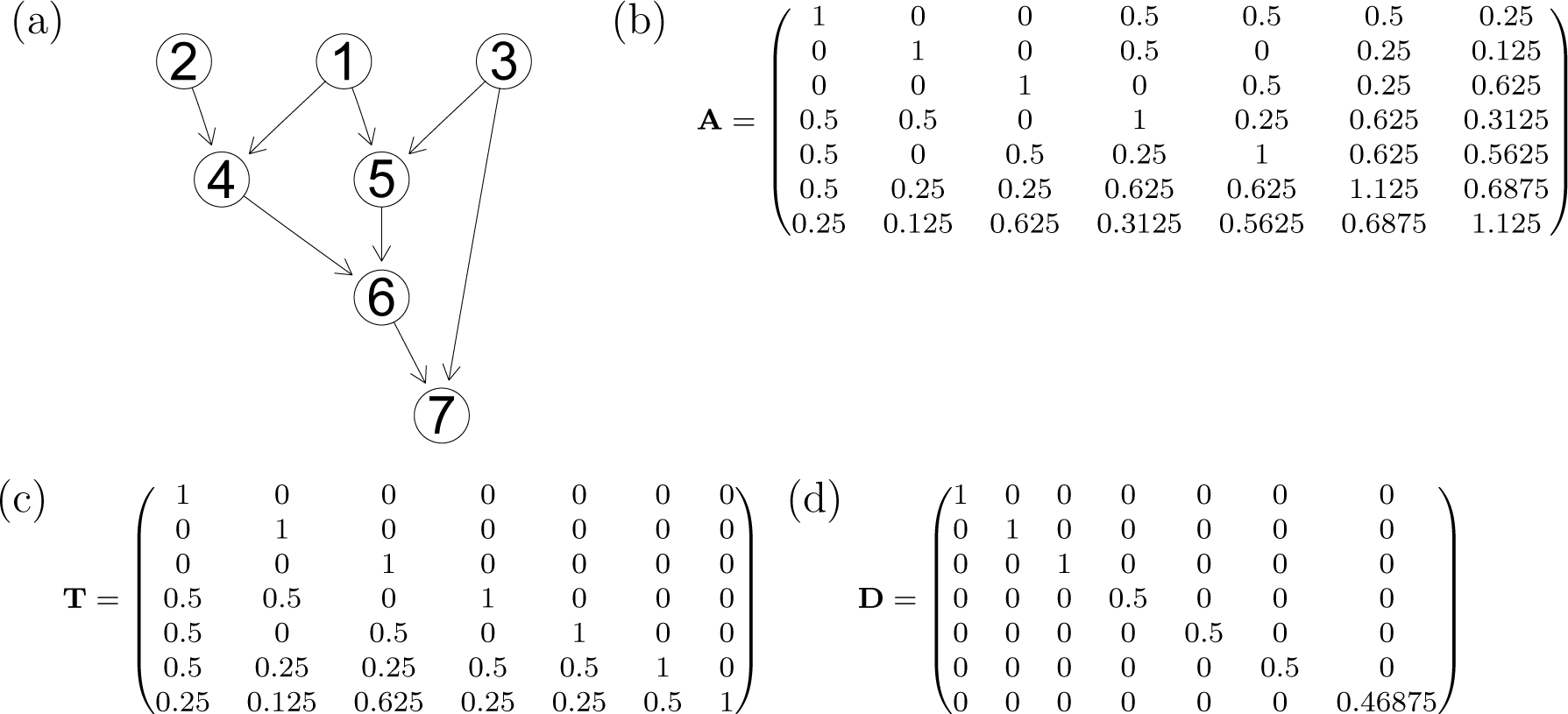
Graphical representation of the pedigree example (a) with the corresponding relatedness matrix **A** (b). The matrices from the decomposition **A** = **TDT**′ are given in (c) and (d). In the genetic group example, animals 1 and 2 are founders of genetic group 1, and animal 3 of genetic group 2.

On the other hand, the diagonal entry *d*_*ii*_ for animal *i* in **D** is calculated as

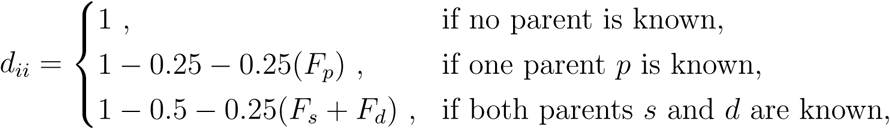

where *F*_*p*_, *F*_*s*_ and *F*_*d*_ are the pedigree-based inbreeding coefficients of the known parent(s) (Mrode, 2005, p. 28). We do not index the inbreeding coefficients with the animal identity (*i*) purely for notational simplicity. For later use we note that, in the absence of inbreeding, the diagonal entry is (1 − 0.5*p*_*i*_), with *p*_*i*_ corresponding to the proportion of *i*’s ancestral genome that is known. Possible values are *p*_*i*_ = 0, 0.5 or 1 if no, one or two parents are known, respectively. This can be understood as follows: If, for example, one parent of an animal *i* is unknown, its predicted breeding value is 0.5 times the breeding value of the known parent, but the other half of its breeding value is unknown. The deviation from the predicted breeding value that could be obtained if both parents were known is absorbed by the Mendelian sampling deviation. The respective variance thus contains the Mendelian sampling variance *plus* a variance that is due to the unknown parent (Kennedy *et al.*, 1988). The more parents are unknown, the larger is this variance.

#### 3.3.2 T and D for genetic groups

In the presence of genetic groups, each unknown parent of an observed animal is assigned to one of the groups, and expected proportions of individual’s genomes that originate from the respective genetic groups can be calculated from the pedigree (Quaas & Pollak, 1981; Wolak & Reid, 2017). For simplicity, we again consider the case with two groups, *g*_1_ and *g*_2_, and denote by **A**_1_ and **A**_2_ the respective relatedness matrices. These can be decomposed in the same way as **A** into

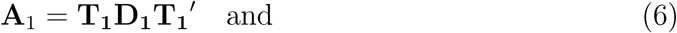

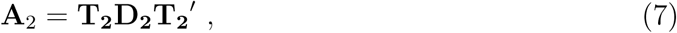

with matrices **T**_1_ and **T**_2_ describing the transmission of alleles through the generations, and Mendelian sampling variance matrices **D**_1_ and **D**_2_. The generalization to more than two groups is straightforward.

Let us assume in the pedigree example of Figure 1 that the parents of founder animals 1 and 2 belong to genetic group 1, and the parents of animal 3 to genetic group 2. This leads to proportional contributions of each genetic group to the genomes of the descending individuals as given by the matrix **Q** (Figure 2a), with columns ***q***_1_ and ***q***_2_ that contain the respective proportions of genetic origin from groups 1 and 2 for each individual. The transmission of alleles within each group is represented by the matrices **T**_*j*_ (*j* = 1, 2). They are designed such that animals with a certain proportion of genetic origin can only pass on the respective fraction of alleles. This means, for example, that an animal *i* with *q*_*i*1_ = 0.5 passes only a proportion of 0.25 (and not 0.5) of alleles to its offspring as part of genetic group 1, while another expected proportion of 0.25 is passed on to its offspring within group 2. The matrices **T**_*j*_ are thus obtained by scaling the respective entries in **T** by the respective group-proportions. This is achieved by multiplying each row of **T** by ***q***_*j*_ or, equivalently, by

**Figure 2:**
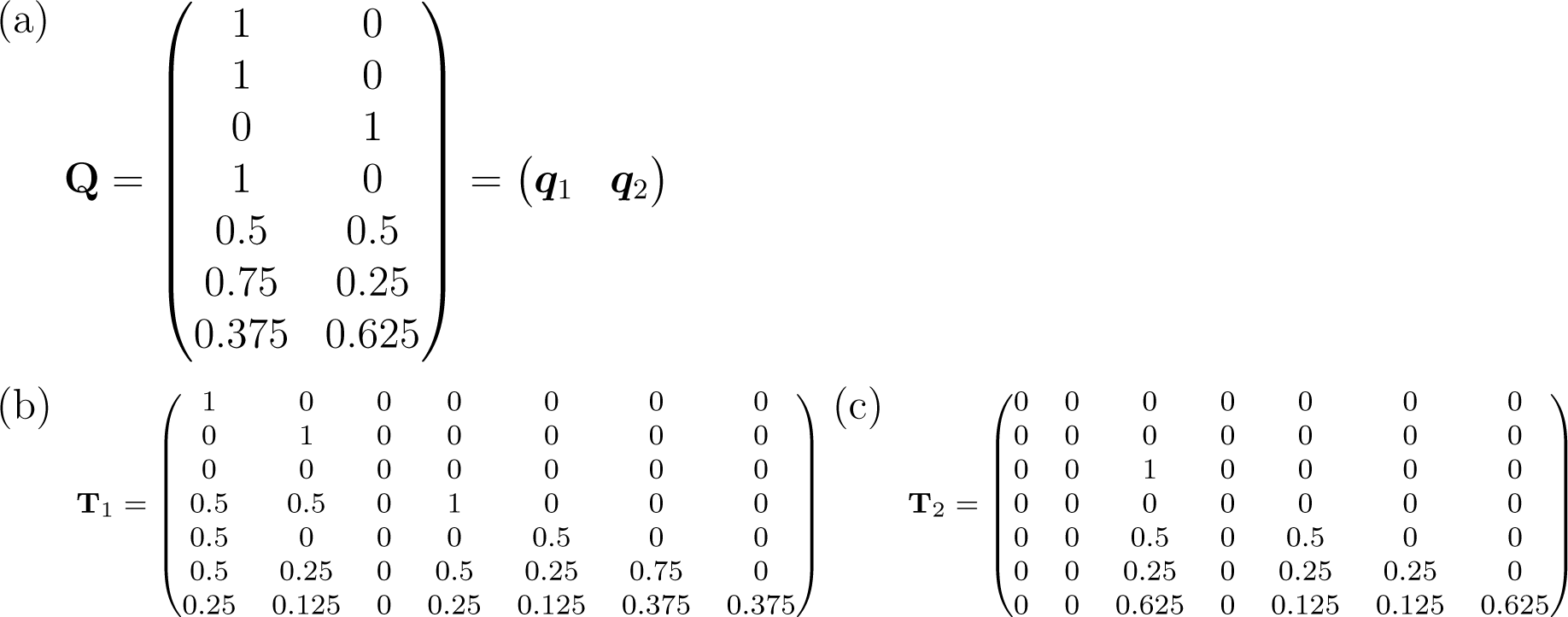
Genetic group matrix **Q** and **T**_*j*_ matrices for the pedigree example. Group-specific proportions of the genome are stored in the **Q** matrix (a). Its columns can be used to derive the group-wise matrices **T**_**1**_ (b) and **T**_2_ (c) in our example.

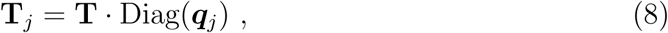

where Diag(***q***_*j*_) denotes a diagonal matrix with diagonal equal to ***q***_*j*_. The matrices **T**_1_ and **T**_2_ for our example are given in Figures 2b and c. Note that the diagonal of **T**_*j*_ corresponds to ***q***_*j*_, which is the respective expected fraction of the genome that belongs to group *j*, and all entries in the respective column are scaled with that same value. Animal 3, for example, that belongs exclusively to group 2 (thus *q*_13_ = 0), has only 0 entries in the third column of **T**_1_, because it cannot transmit any group-1-alleles to its descendants. Animal 5, on the other hand, has *q*_15_ = 0.5, thus the fifth column in **T** is multiplied by 0.5 to obtain the respective column in **T**_1_.

Next, we need to find appropriate versions of **D**_1_ and **D**_2_. We noted in Section 3.3.1 that, in the absence of inbreeding, *d*_*ii*_ = 1 − 0.5*p*_*i*_ with *p*_*i*_ representing the proportion of the ancestral genome that is known. To calculate the respective entries 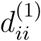 and 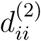 in the group-specific matrices, we have to multiply *p*_*i*_ by the proportions of genetic origin *q*_*i*1_ and *q*_*i*2_, because the respective product then corresponds to the ancestral proportions that are known *within* the respective group. In the case where only one parent is known, multiplication must be with the genetic proportion of the known parent, denoted here as 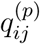, because only this respective part of the ancestral genome within group *j* is then known. This leads to

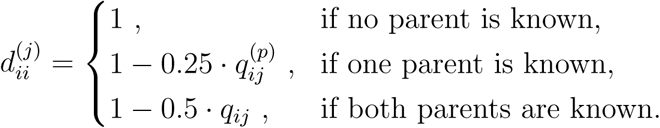

Let us now also account for inbreeding, which may influence 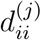 for individuals with at least one parent known. Starting with animals that have both parents known, and rearranging the entries *d*_*ii*_ = 1 − 0.5 − 0.25(*F*_*s*_ + *F*_*d*_) in the original matrix **D**, leads to

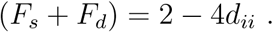

The knowledge of (*F*_*s*_ + *F*_*d*_) for each animal is useful to derive the entries 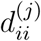 for group *j* in the presence of inbreeding: By scaling the effect of inbreeding with the group-specific proportions *q*_*ij*_ of an animal’s genome, the entries are given as

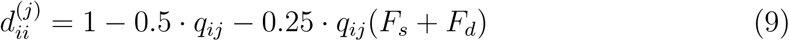

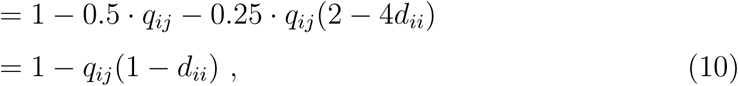

where the third line is an algebraic simplification of the second line. The same calculation for animals with only one parent known, using *d*_*ii*_ = 1 − 0.25 − 0.25(*F*_*p*_) and solving for *F*_*p*_, leads to the same formula with *q*_*ij*_ replaced by 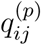 from the known parent. Finally, if both parents are unknown (*i. e*. for *d*_*ii*_ = 1), the formula also leads to the correct value of 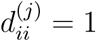. Applying formula (10) to the above pedigree example leads to **D**_1_ and **D**_2_ (Figures 3a and b).

**Figure 3:**
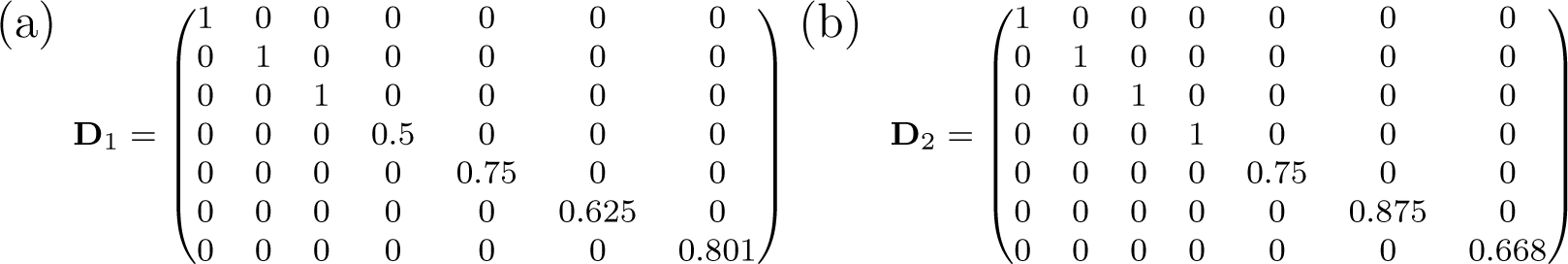
Group-specific matrices **D**_1_ and **D**_2_ for the example pedigree.

Formula (10) is simple and convenient. However, it provides only an approximation of the correct matrix entries, because in (9) we assumed that parental inbreeding can simply be scaled by the genetic group proportions *q*_*ij*_ (for two known parents) or 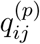 (for one known parent). Instead, the theoretically correct way to deal with parental inbreeding coefficients to derive 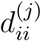 would be to use the actual *partial* (*i. e*. group-specific) parental inbreeding coefficients, denoted *e. g*. as 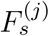 or 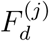 for parents *s* and *d*. These group-specific inbreeding coefficients contain only the inbreeding that emerge due to inbreeding *within genetic group j*, that is, they measure the probability that an individual is identical by descent for an allele that descended from founders within group *j* (Lacy *et al.*, 1996). The correct way to calculate 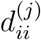 is thus given by

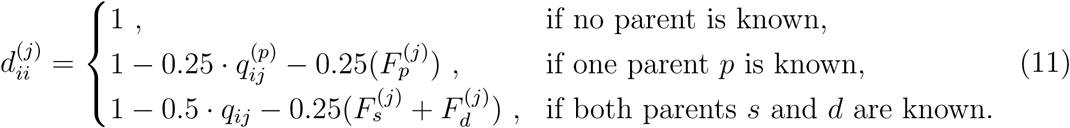

Obviously, this formula requires the calculation of group-specific inbreeding coefficients, which is computationally cumbersome. One way to obtain these coefficients is by first calculating *founder*-specific inbreeding coefficients that partition the total inbreeding coefficient *F*_*i*_ into the additive components *F*_*ik*_ from each founder animal *k*, as proposed by Lacy *et al.* (1996). Because partial contributions for all founders sum up to *F*_*i*_ (*e. g*. Lacy *et al.*, 1996; Gulisija *et al.*, 2006), we can sum only over founders from genetic group *j* to obtain group-specific inbreeding coefficients 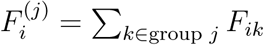.

Let us illustrate the difference between the approximate method suggested in equation (10) and the correct formula for 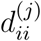 given in (11) for our example from Figure 1. Animal 6 is the only parent in the pedigree with a non-zero inbreeding coefficient, which is *F*_6_ = 0.125. However, because animals 1 and 2 are founders of group 1 and animal 3 is a founder of group 2, the pedigree reveals that inbreeding originates only from matings within group 1. Therefore, *F*_6_ is split into group-specific inbreeding coefficients as 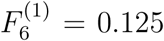 and 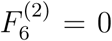. By plugging these values into (11) to estimate the respective values for animal 7 (which is the only animal that is affected by this change), we obtain 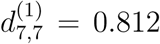 and 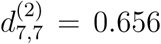, which are quite close to the approximate values 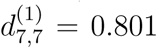 and 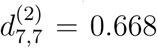 from Figures 3a and b. Note that in this paper we will continue to use the convenient and computationally efficient approximation (10) to scale the entries in **D**_*j*_, but we will illustrate the consequences of this approximation with simulations and for our application to the house sparrow example below and in Appendix 1.

#### 3.3.3 Properties of group-specific relatedness matrices

Once the components **T**_*j*_ and **D**_*j*_ for each group *j* are known, a simple matrix multiplication yields the group-specific relatedness matrices 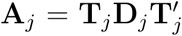. These are given in Figure 4 for the two groups considered in our example. An important aspect is that both **A**_1_ and **A**_2_ contain columns and rows with all variances and covariances equal to zero, namely for animals *i* without a contribution from group *j* (*q*_*ij*_ = 0). While this is theoretically correct, because the respective breeding value is then *a*_*ij*_ = 0, the resulting matrices are singular. When it comes to implementation, the problem can be solved by replacing zeros on the diagonal by very small values, for example 10^-6^ or even 10^-12^. The choice is not critical in our experience, but it would be prudent to check the robustness of the results to different values.

**Figure 4:**
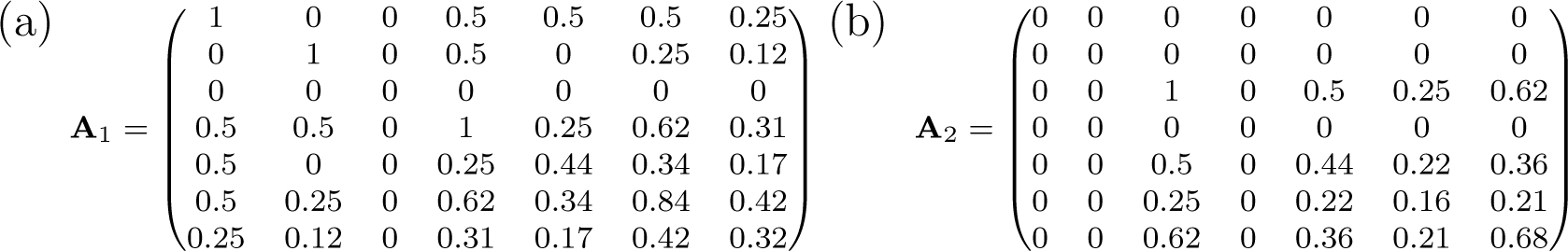
Group-specific relatedness matrices for the example pedigree with entries rounded to two digits.

#### 3.3.4 Scaling the inverse relatedness matrix

In practice it is usually the inverse relatedness matrix **A**^-1^, not **A**, that is calculated and stored directly from the pedigree. Consequently, it is computationally more convenient, and in most cases much more efficient, to apply the appropriate scaling and transformation operations directly to **A**^-1^ to obtain group-specific inverses 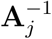. From (5) it follows that the inverse of **A** can be decomposed into

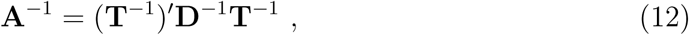

where **D**^-1^ = Diag(1/*d*_11_*, …,* 1/*d*_*nn*_). Using that **T**_*j*_ = **T** *·* Diag(***q***_*j*_), and the decomposition of **A**_*j*_ as given *e. g*. in equation (6), some matrix algebra shows that

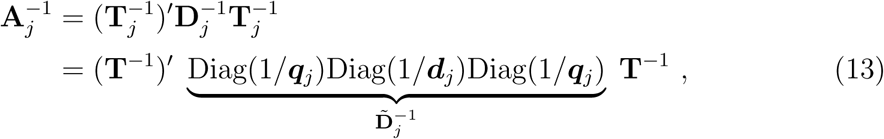

where 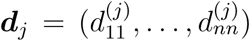 are the diagonal entries of **D** _*j*_ as derived from equation (10), and ***q***_*j*_ is again the vector of genetic proportions inherited from group *j*. This illustrates that it is sufficient to calculate the (computationally relatively expensive) generalized Cholesky decomposition **A**^-1^ into **T**^-1^ and **D**^-1^ only once, and to derive 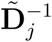 with diagonal entries 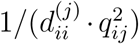 to obtain 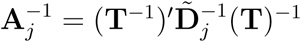 as in (13) for each group. Again, entries with *q*_*ij*_ = 0 are replaced by very small values, *e. g*. 10^-12^, to avoid singularities.

## 4 Simulation

### 4.1 Generating data

To illustrate the performance of genetic group models with group-specific additive genetic variances we simulated data using the `simGG()` function from the R package `nadiv` (Wolak, 2012). The function allows generating pedigrees and phenotypes for a focal population (group 1) that receives a specified number of immigrants from another population in each generation (group 2). Group-specific mean breeding values and additive genetic variances can be determined by the user, and breeding values for the founder animals of both genetic groups are sampled from the respective distributions. Offspring breeding values are calculated from the parental mean, plus a Mendelian sampling deviation that depends on the additive genetic variance of the resident population, but there is no additional term that introduces a segregation variance (for more details, see Wolak, 2012). The simulation assumes random mating among individuals that currently live in the same population, thus offspring may inherit genetic components from both genetic groups due to immigration. The contributions *q*_*ij*_ from group *j* for animal *i* were calculated with the `ggcontrib()` function from the `nadiv` package. For data generated with group-specific mean breeding values and additive genetic variances, the appropriate underlying model for the analysis is

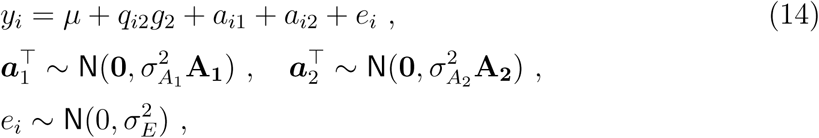

where we used the same notation as in Section 3, and fixed the focal group mean *g*_1_ = 0 for identifiability reasons (thus the term *q*_*i*1_*g*_1_ is omitted from equation (14)). We simulated data according to three different scenarios, but always setting the population mean *µ* = 10, group means *g*_1_ = 0, *g*_2_ = 2, group-specific variances 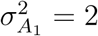 and 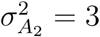 and residual variance 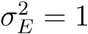. Each scenario encompassed 10 non-overlapping generations.

**Scenario 1** The carrying capacity of the population was set to 300 individuals. In each generation, 100 mating pairs were created by random sampling with replacement from the adults, and each pair contributed 4 offspring to the next generation. In addition, 30 immigrants were added to the population in each (except the first) generation. Finally, a subset of the offspring was randomly selected such that the population size always corresponded exactly to its carrying capacity.

**Scenario 2** This scenario was the same as scenario 1, except that only 5 (instead of 30) immigrants were allowed in each of the non-overlapping generations, so that animals of the immigrant group were rare.

**Scenario 3** In this scenario we used the same carrying capacity as above, 20 immigrants per generation, but we only allowed for 5 breeding pairs per generation that produced 60 offspring each. While this scenario has an immigration rate that lies between scenarios 1 and 2, the low number of breeding pairs induces higher inbreeding levels. This scenario is therefore suitable to illustrate the consequence of using approximation (10) to scale the group-specific Mendelian sampling variance matrices **D**_*j*_, which is only (potentially) critical in the presence of inbreeding, because the approximation affects only the scaling of parental inbreeding coefficients.

For each scenario, we generated 100 datasets, and each of them was analyzed with the genetic group animal model that accounted for heterogeneous additive genetic variances 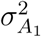 and 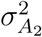, as given in (14), and compared to the outcome of the standard genetic groups model that allowed for different mean breeding values, but only a single (homogeneous) variance in both groups, given as

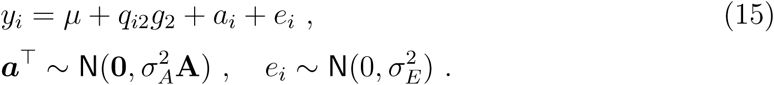

For scenario 3 we also investigated how close the group-specific Mendelian sampling variance approximations for 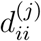 from equation (10) are in comparison to the correct version given in (11). The group-specific inbreeding coefficients present in the correct formula were calculated with the software `GRain` (Baumung *et al.*, 2015, details are given in Appendix 1). Correlation coefficients *ρ* between the (correct) 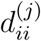 values from (11) and the approximated values from formula (10) were calculated, and all simulations were analyzed with both versions for comparison.

Following the recommendation by He & Hodges (2008), we stored posterior modes (and not posterior means) of the variance components in each iteration. All models were fitted with integrated nested Laplace approximations (INLA, version from June 20, 2017), which provides a fast and accurate alternative to MCMC (Rue *et al.*, 2009), although it has so far only rarely been used for animals models (but see *e. g*. Holand *et al.*, 2013; Steinsland *et al.*, 2014; Roulin & Jensen, 2015). All variance components were given penalized complexity (PC) priors, which were suggested as valid alternatives to the (less recommended) gamma priors (Simpson *et al.*, 2017). The PC(*u, α*) prior has an intuitive parameterization: The prior probability for the standard deviation *σ* is given as Pr(*σ* > *u*) = *α* (with 0 < *α* < 1). Here we used PC(1, 0.05) priors for all variances (thus Pr(*σ* > 1) = 0.05 *a priori*), but results were insensitive to this choice. All fixed effect parameters were assigned independent **N**(0, 10^4^) priors. A short tutorial including R code to generate and analyze data for the models used here can be found in Appendix 4.

### 4.2 Simulation results

Using the appropriate model (14) with heterogeneous group variances resulted in estimates that were close to the variances used to generate the data in scenarios 1 and 3, while the model estimates in scenario 2 suffer from large uncertainty (Figure 5, left). In particular the variance 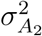 of the underrepresented immigrant population in scenario 2 was difficult to identify and biased towards 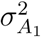 (Figure 5c). The results thus indicate that the genetic group model (14) is able to isolate approximately correct group-specific additive genetic variances, but that some caution is required if representatives of a genetic group are rare in the data set: Group-specific additive genetic variances are then not completely identifiable.

**Figure 5:**
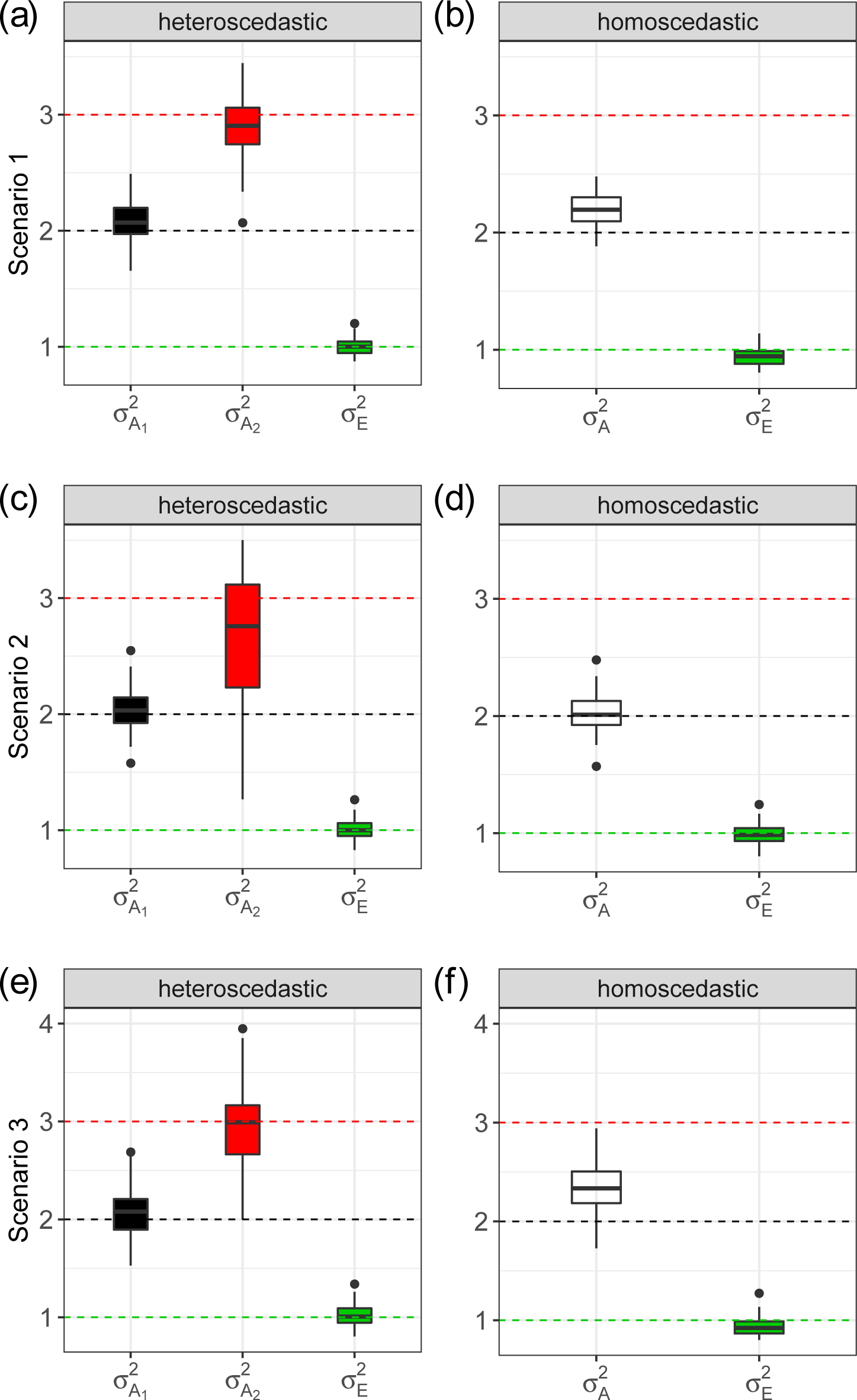
Results from 100 iterations for simulation scenarios 1–3. The boxplots represent the distributions of estimated variances (posterior modes) from a model with genetic groups and heteroscedastic additive genetic variances 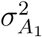 and 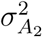 (left panel), compared to the results from a model that only allowed for a single homogeneous variance 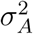 (right panel). Dashed lines indicate the references that were used to generate the data (black: 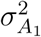, red: 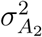, green: 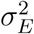).

When the simulated data were fitted using the genetic group model with a single, homoscedastic variance 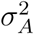 (equation (15)), the resulting estimates were generally between the two simulated group-specific variances (Figure 5, right). In the presence of only few immigrants, the estimate tended to be close to 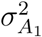 (scenario 2), while it tended towards 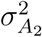 when there was more immigration (scenarios 1 and 3). This is as expected, and illustrates that genetic group models with a single, homogeneous variance will estimate a value in between the true group-specific additive genetic variances, with a tendency towards the variance of the more numerous group. These patterns were qualitatively similar when the additive genetic variances of the immigrant and resident population were switched, such that residents had larger additive genetic variance than immigrants (results not shown).

Simulation scenario 3 led to datasets with a mean inbreeding coefficient of *F* = 0.10. Interestingly, the comparison between the approximate versus the correct values in the **D**_1_ and **D**_2_ matrices shows that the approximation suggested in equation (10) leads to 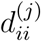 values that are highly correlated with the correct values from equation (11). As an example, we found correlation coefficients *ρ* ≥ 0.988 in three randomly selected simulation runs. In addition, using the correct **D**_*j*_ matrices led to distributions of estimated variances that are indistinguishable from the results when the approximations were used (see Figures S1 and S2 in Appendix 1).

## 5 Application to house sparrow data

### 5.1 Study population

As a proof of concept, we applied our method to empirical data from a long-term study of an insular house sparrow meta-population off the Helgeland coast in northern Norway. The study has been running continuously since 1993, and is used as a model system to examine ecological and evolutionary processes in fragmented vertebrate populations (*e. g*. Ringsby *et al.*, 2002; Jensen *et al.*, 2008; Pärn *et al.*, 2012; Baalsrud *et al.*, 2014). The islands in the meta-population differ in characteristics related to environmental conditions, habitat type and population size, with considerably larger and more stable populations on the five islands that are located closer to the mainland (denoted as *inner* islands) compared to the three islands located further away (denoted as *outer* islands). The ten remaining islands are summarized as *other* islands (see Figure S4 in Appendix 2 for an overview of the island system).

Small blood samples were collected from all captured birds on the eight inner and outer islands to provide DNA for Single Nucleotide Polymorphism (SNP) genotyping on a 200K SNP array (see Lundregan *et al.*, 2018). Only successfully genotyped birds that were also measured for body mass and/or wing length were included in our animal model analyses. For inner and other islands, the dataset included phenotypic measurements taken during the breeding seasons since 1993 and 1995, respectively. Due to strong population bottlenecks on the outer islands in 2000 (Baalsrud *et al.*, 2014), only measurements taken since 2002 were used for the populations in the outer group. Details on how morphological measurements of wing length and body mass were taken on adult birds are given by *e. g*. Jensen *et al.* (2004, 2008).

Parentage analyses for the eight island populations in this study were carried out with the R package `SEQUOIA` (Huisman, 2017). Briefly, SNP genotype data of all adults recorded as present on any of the eight inner or outer islands during the years 1998-2013 (the inner group) or 2003-2013 (the outer group) were used in the parentage analyses. This resulted in a “meta-population pedigree” (*N* = 3116) spanning up to 14 generations, where both parents were known for 52.7%, one parent was known for 25.0%, and no parents were known for 22.3% of the individuals. Since `SEQUOIA` introduces *dummy parents* to preserve known relationships, *e. g*. sibling relationships, even when parents are not genotyped, a higher percentage of individuals had “known” parentages (81.0%, 5.5% and 13.5% with two, one or no parents known, respectively).

The genetic group analysis that we carry out here requires that each unknown parent (*i. e*. each founder) must be assigned to one of the genetic groups. That is, we must attribute unknown parents to the *inner, outer* or *other* island group to determine their genetic origin. This was done here by first identifying the natal (hatch) island of all individuals, either from ecological data or, if unavailable, by using genetic assignment procedures based on the SNP genotype data (see Saatoglu *et al.*, 2018). Of all individuals in our dataset, 1436, 481, and 64 individuals were assigned to a natal island in the inner, outer and other group, respectively (Saatoglu *et al.*, 2018). This information was then used to assign missing parents to the genetic group to which the hatch island of the respective individual belonged. As an example, if an individual with missing parents is known to be born on one of the inner islands, the respective missing parents were assumed to be founder individuals of the inner island group, and so on. Finally, because inbreeding depression is known to occur in our study system (Jensen *et al.*, 2007; Billing *et al.*, 2012), we accounted for any inbreeding effects on body mass and wing length (Reid *et al.*, 2006; Reid & Keller, 2010) by including each individual’s genomic inbreeding coefficient *F*_GRM_ (Yang *et al.*, 2011a) as a covariate in all models fitted here, where *F*_GRM_ was estimated as described by Niskanen *et al.* (2018).

### 5.2 Analysis of wing length and body mass

With all unknown parents assigned to one of the genetic groups (*inner, outer, other*), each individual *i* obtained expected proportions of genetic origin *q*_*ij*_ from the three groups *j* by propagating the founder individual’s genome through the pedigree using the `ggcontrib()` function. Note that the different groups are very unequal in sampling size, which can be seen from summation of genetic proportions over all animals within a group *j*, that is, *n*_*j*_ = Σ_*i*_ *q*_*ij*_, which corresponds to equivalents of full animal genomes (Table 2). This is not surprising given the smaller population sizes (Baalsrud *et al.*, 2014), lower recapture rates (Holand *et al.*, 2016), and shorter time-series for populations in the outer islands group, and considering that there were no genotyping efforts on the other islands, thus we only see genome from the other islands if it was introduced via an immigration event to one of the inner or outer islands.

**Table 2:** Estimates (posterior modes) and 95% CIs (in brackets) of the three group-specific additive genetic variances 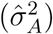 for inner, outer and other genetic groups, as well as for a single homogeneous variance across groups for the two traits body mass and wing length. The sample sizes denote the equivalent of full animal genomes that are present in the three genetic groups (*n*_*j*_, for the model with heterogeneous variances) or in the total dataset for the respective trait (*n*, for the homogeneous model). For comparison, the phenotypic variances 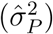 in the three groups and for the total population, calculated only from the 1113, 148 and 33 pure-bred animals for mass and from the 1117, 146 and 33 pure-bred animals for wing length in the inner, outer and other groups.

|  | Body mass |  |  | Wing length |  |  |
| --- | --- | --- | --- | --- | --- | --- |
| | $\hat{\sigma}_A^2$ | $n_j$ or $n$ | $\hat{\sigma}_P^2$ | $\hat{\sigma}_A^2$ | $n_j$ or $n$ | $\hat{\sigma}_P^2$ |
| *inner* | 1.56 (1.15, 2.06) | 1572.5 | 5.72 | 1.77 (1.53, 2.13) | 1571.0 | 5.34 |
| *outer* | 0.93 (0.42, 2.24) | 267.3 | 5.65 | 2.29 (1.78, 3.44) | 262.6 | 6.48 |
| *other* | 0.70 (0.26, 2.85) | 131.3 | 4.87 | 1.00 (0.59, 2.69) | 129.4 | 6.02 |
| *total* | 1.54 (1.19, 2.02) | 1971 | 5.53 | 1.79 (1.60, 2.21) | 1963 | 5.41 |

For the two traits investigated here, mass (in g) and wing length (in mm), we fitted separate models that accounted for sex (0=males, 1=females), inbreeding *F*_GRM_, month of measurement (numeric with values 5, 6, 7, and 8) and age as fixed effects that were stored in matrix ***X***, and current island of residence where the measurement was taken (*island*), hatchyear (*year*), animal (*id*) and an independent residual term (*e*) as random factors. An individual *i* was included in the model if it had at least one recorded observation *k* of the respective trait. The model with group-specific mean and variances of the breeding values is thus given as

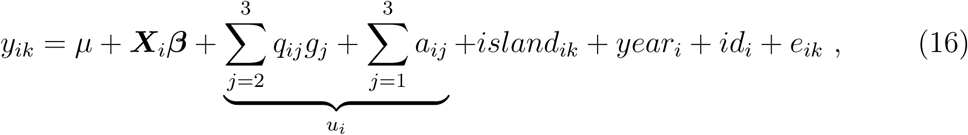

where the total genetic contribution *u*_*i*_ is the sum of the weighted means *g*_*j*_ and the variability from the different genetic groups, as introduced in equation (4). The three genetic groups *inner, outer* and *other* are encoded as groups 1, 2 and 3, respectively, where the mean of the inner group was set to *g*_1_ = 0 for identifiability reasons. The estimates *g*_2_ and *g*_3_ thus reflect differences in group means with respect to the inner population. The components *a*_*i*1_, *a*_*i*2_ and *a*_*i*3_ are distributed with mean zero and heterogeneous additive genetic variances 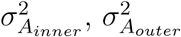, and 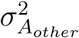, with dependency structures given by the group-specific relatedness matrices **A**_*inner*_, **A**_*outer*_ and **A**_*other*_ that were calculated as explained in Section 3. The results from fitting model (16) were compared to the standard genetic group model that only accounts for differences in group means, but with homogeneous additive genetic variance 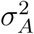 and dependency structure defined through the relatedness matrix **A**. Both models were fitted to the data using INLA. All variances were given PC(1, 0.05) priors, and fixed effects parameters were assigned independent **N**(0, 10^4^) priors.

Interestingly, the results indicate that the outer group has a somewhat smaller estimated additive genetic variance than the inner and other groups in the case of body mass (Table 2, left), while the estimated additive genetic variance is larger for the outer group for wing length (Table 2, right), although for both traits the respective 95% credible intervals (CI) are relatively large and overlap. The differences in additive genetic variance estimates could indicate that animals living on the three island groups show different levels of variation in the genes that affect the phenotypic traits, *e. g*. due to different allele frequencies, although we cannot rule out that additional shared environmental effects are confounded with additive genetic variances (Stopher *et al.*, 2012). In both cases, the variance estimate 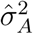 from the homogeneous model lies between the group-specific variance estimates, with a tendency towards the estimates for the inner population, which is by far the largest group. These results were also compared to the observed phenotypic variance 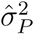 of the two traits, where the respective group-specific phenotypic variances were only calculated using the pure-bred animals (*i. e*. those with *q*_*ij*_ = 1) in each group (Table 2). It is also interesting to note that the estimates for *g*_2_ indicate that animals on the outer islands are lighter and have somewhat shorter wings than animals on the inner islands (Table 3), which is in agreement with earlier findings (Jensen *et al.*, 2004, 2008; Holand *et al.*, 2011). The remaining variance estimates of the model are not of primary interest and are thus given in Table S3 of Appendix 2.

**Table 3:** Posterior means and 95% CIs of the fixed effects for the animal models used for body mass and wing length. The estimates were extracted from models with either group-specific (heterogeneous) additive genetic variances, or a single (homogeneous) additive genetic variance.

|  | Body mass |  | Wing length |  |
| --- | --- | --- | --- | --- |
|  | heterogeneous | homogeneous | heterogeneous | homogeneous |
| sex | 0.46<br>(0.29, 0.64) | 0.48<br>(0.30, 0.65) | -2.76<br>(-2.89, -2.63) | -2.77<br>(-2.90, -2.64) |
| $F_{GRM}$ | -1.23<br>(-3.09, 0.63) | -1.24<br>(-3.09, 0.61) | -1.32<br>(-2.69, 0.06) | -1.36<br>(-2.74, 0.01) |
| month | -0.29<br>(-0.35, -0.24) | -0.30<br>(-0.36, -0.24) | -0.18<br>(-0.22, -0.15) | -0.19<br>(-0.22, -0.15) |
| age | 0.08<br>(0.02, 0.14) | 0.08<br>(0.02, 0.14) | 0.47<br>(0.43, 0.50) | 0.47<br>(0.43, 0.50) |
| $g_2$ (outer) | -0.83<br>(-1.29, -0.39) | -0.84<br>(-1.38, -0.29) | -0.44<br>(-0.82, -0.01) | -0.62<br>(-1.09, -0.12) |
| $g_3$ (other) | -0.28<br>(-0.84, 0.27) | -0.18<br>(-0.85, 0.50) | 0.08<br>(-0.34, 0.49) | -0.07<br>(-0.64, 0.51) |

All results presented here involved the approximate approach from formula (10) to scale the group-specific **D**_*j*_ matrices. To illustrate that this approximation is unproblematic, we repeated all calculations with the correct versions as given in equation (11), which again involved the gene dropping method provided in `GRain`. Details are given in Appendix 1, and all results remain essentially unchanged (see Figure S3 and Tables S1 and S2 in Appendix 1). Finally, it is worth reiterating that model (16) does not account for the three segregation variances that would occur between any two groups, because these are expected to be negligibly small. In addition, estimating these three additional variances would impose unrealistic requirements on these data. To illustrate that ignoring these variances is not critical, we also fitted a model with a segregation term for the sparrow example with only two genetic groups (inner and outer). Details on how to estimate segregation variances are given in Appendix 3. Importantly, the results in Table S4 confirm that segregation terms are indeed very small, and that their inclusion only leads to irrelevant changes in the results.

## 6 Discussion

We have introduced an extension of the animal model that allows for unequal additive genetic variances in the presence of multiple interbreeding genetic groups. Our method requires that group-specific relatedness matrices **A**_*j*_ are derived. To this end, the full relatedness matrix **A** is decomposed into matrices **T** and **D**, and simple algebraic scaling operations are used to derive the group-specific versions **T**_*j*_ and **D**_*j*_ that are then again multiplied to obtain **A**_*j*_. The method is computationally efficient, in particular when an (accurate) approximation for the group-specific Mendelian sampling variance matrices **D**_*j*_ is used. Although genetic group animal models have been used before, in particular in animal and plant breeding setups, modeling heterogeneous additive genetic variances has so far been considered unfeasible (Wolak & Reid, 2017). Therefore, natural populations have only been analyzed with genetic group models that account for mean differences in additive genetic effects, but not for heterogeneous variances. Here we overcome this limitation by assuming that segregation variances are often negligible for polygenic traits. How often will this assumption hold? The segregation variance from formula (3) approximates zero whenever the *infinitesimal model*, which underlies the animal model and which posits that traits are determined by a large number of genes with small effects, holds approximately. This is likely the case for many polygenic, complex traits (Barton *et al.*, 2017). Our estimates of the segregation variance in the empirical house sparrow data set support the view that segregation variance may often be negligible. This result is mirrored in GWAS of the genetic architectures of body mass and wing length in house sparrows and other passerines, which revealed a polygenic basis for these traits, where any significant genomic region explains only a very small proportion of the phenotypic variance (Santure *et al.*, 2015; Silva *et al.*, 2017). Taken together, these results suggest that segregation variances can often be neglected in genetic group models, provided the traits of interest are truly very polygenic. When focal traits have a genetic architecture with only few causal genes with a large effect, omitting the segregational variance may however introduce a non-negligible bias in the estimated additive genetic variances. In such a case it is still possible to formulate a model that accounts for segregation variances, as explained in Appendix 3, although such models may quickly impose unrealistic demands on the data.

Estimating and disentangling variance components is generally known to be difficult, and it is particularly challenging for genetic group models with group-specific additive genetic variances. The problem is that the group-specific covariance matrices **A**_*j*_ are the sole sources of information that allow to discriminate the variance components, yet these matrices may be similar in the presence of many multi-bred animals. The results from the house sparrow example in Table 2 illustrate that group-specific variance estimates may suffer from larger uncertainty than a single homogeneous variance, especially when group sizes are small, as it is the case for the *outer* and *other* groups. Allowing for heterogeneous variances is therefore only advocated if there is an actual need for them. While this is often not evident in advance, the user may want to fit the homogeneous and the heterogeneous models to determine if the more complex model is needed. A possible “objective” way to find this out might be via the use of information criteria, such as the deviance information criterion (DIC) for Bayesian models (Spiegelhalter *et al.*, 2002), although we do not generally recommend to rely on it (for some DIC criticism see *e. g*. Plummer, 2008; Hodges, 2014). Interestingly, in the sparrow example the DIC for the heterogeneous model for mass was 4 units larger than for the model with a single variance, while it was −15 units smaller for wing length. Thus there is some weak indication that the more complex model is not needed when body mass is analyzed, while there appears to be some improvement for wing length, which is consistent with the observation that the estimated variances show less overall differences for body mass than for wing length (Table 2). In principle, the variance estimates can also be used to assess group-specific versions of heritability (Falconer & Mackay, 1996) or evolvability (Houle, 1992), where the additive genetic variance is replaced by the group-specific version 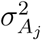.

A general limitation of genetic group models is that parent-offspring relations are needed to propagate genetic contributions from founder individuals through the pedigree to determine **Q**. Genomic relatedness matrices alone are therefore not sufficient to fit genetic group models, but a combination of genomic and pedigree information, the latter possibly inferred from genetic data (Ko & Nielsen, 2017), may provide a powerful basis for genetic group models.

The proposed extension of genetic group models will be useful for any study population that is structured into subpopulations, given that sufficient information on migration or crossbreeding events is available. In particular, the fact that group-specific additive genetic variances can be estimated for subpopulations that are not completely isolated might also be useful when interest centers around the dependency of additive genetic variance on the effective population size, a relation that is of pivotal interest in evolutionary and conservation biology. Finally, the method may provide a starting point for the estimation of temporal or spatial variation of additive genetic variance.

## Acknowledgements

We are grateful to Dr. Timothée Bonnet for comments to the manuscript. This work was mainly funded by the Faculty of Science of the University of Zurich (S.M., L.F.K.). We thank the many researchers, students, and fieldworkers for their contributions to collecting the empirical data on house sparrows, and laboratory technicians for assistance with laboratory analyses. This study was supported by grants from the Norwegian Research Council (programmes STORFORSK, Strategic University Program in Conservation Biology, project 221956 to H.J.), the Norwegian Directorate for Nature Management, the EU-commission (project METABIRD), and the Academy of Finland (project 295204 to A.N.). This work was also partly supported by the Research Council of Norway through its Centres of Excellence funding scheme, project 223257).

